# Contribution of cellular macromolecules to the diffusion of a 40 nm particle in *Escherichia coli*

**DOI:** 10.1101/2024.01.22.576611

**Authors:** José Losa, Matthias Heinemann

## Abstract

Due to the high concentration of proteins, nucleic acids and other macromolecules, the bacterial cytoplasm is typically described as a crowded environment. However, the extent to which each of these macromolecules individually affects the mobility of macromolecular complexes, and how this depends on growth conditions, is presently unclear. In this study, we sought to quantify the crowding experienced by an exogenous 40 nm fluorescent particle in the cytoplasm of *E. coli* under different growth conditions. By performing single particle tracking measurements in cells selectively depleted of DNA and/or mRNA, we determined the contribution to crowding of mRNA, DNA and remaining cellular components, i.e., mostly proteins and ribosomes. To estimate this contribution to crowding, we quantified the difference of the particle’s diffusion coefficient in conditions with and without those macromolecules. We found that the contributions of the three classes of components were of comparable magnitude, being largest in the case of proteins and ribosomes. We further found that the contributions of mRNA and DNA to crowding were significantly larger than expected based on their volumetric fractions alone. Finally, we found that the crowding contributions change only slightly with the growth conditions. These results reveal how various cellular components partake in crowding of the cytoplasm and the consequences this has for the mobility of large macromolecular complexes.

**Statement of Significance:** The mobility of a particle of interest in the cytoplasm depends on a variety of factors that include the concentration, shape and physicochemical properties of crowding obstacles. Different macromolecules in the cell are therefore expected to hinder the mobility of a given particle to different extents. However, an accurate and systematic investigation of these hindrances to mobility *in vivo* has not been yet carried out. In this work, through a novel combination of experimental and computational approaches, we determine the diffusion coefficient of a 40 nm particle in the cytoplasm of *E. coli* under conditions of selective removal of some macromolecules. This allows us to quantify the hindering effect of each of the depleted macromolecules on the mobility of the said particle. For DNA, mRNA, and remaining macromolecules, we observe that this effect is of comparable magnitude, being largest in the latter case. This work sheds light on the interplay between intracellular composition and the physical properties of the cytoplasm at the 40 nm scale.

## Introduction

Due to the high concentration of intracellular macromolecules, with proteins, for instance, reaching ∼300 g/L in bacterial cells (1), the interior of a cell is often described as a “crowded” environment (2–4). By taking up space that is otherwise occupied by water or small solutes, macromolecules reduce the effective free intracellular volume, and can thereby increase the chemical activity of cellular components, such as proteins and ribosomal subunits, by several orders of magnitude (5, 6). In addition, by causing proteins to associate into active multimeric complexes, by increasing local reactant concentrations, or by enhancing reaction kinetics, crowding can be beneficial for cellular processes (5–9). On the other hand, high levels of crowding hinder the diffusion of both macromolecules and complexes (e.g., ribosomes and acyl-tRNAs) to such an extent that certain processes become diffusion-limited (9, 10). Crowding is therefore a defining characteristic of the intracellular environment, and a better understanding of how various macromolecules contribute to this effect is needed.

Multiple studies have elucidated how crowding by certain cellular components affects the organization of the intracellular environment. DNA, for instance, has received particular attention. In *Escherichia coli* , the nucleoid has been shown to exclude (poly-)ribosomes (11–14) and other components with dimensions larger than its characteristic “mesh size”, of about 50 nm (15), leading to the accumulation of such components at the polar regions. In turn, the highly abundant ribosomes, which account for approximately 30% of the dry mass of a bacterial cell (ref. 16; see also estimation in Supplemental Note 1.5), are themselves major sources of crowding, and changes in their abundance have recently been shown to regulate liquid-liquid phase separation events in yeast (17).

Following its definition as the fraction of excluded volume in a cell, crowding is a feature that depends on the relative size of a tracer particle compared to that of the crowder macromolecules. Indeed, particles of different sizes do not “experience” the same excluded volume. While small particles can diffuse in- and out of the voids formed between crowder molecules, larger particles are prevented from accessing, i.e., are excluded from, any void that is smaller than their characteristic length (6, 18). Thus, the exclusion of polyribosomes from the nucleoid region (19–21) happens despite the low volumetric fraction of DNA in the cell, of only 1.4% (22). In contrast, smaller sized proteins or ribosomal subunits are unaffected by the presence of the nucleoid and are homogeneously distributed across the cytoplasm, at least under normal growth conditions (12, 20, 23). Because of this length-scale dependence, previously developed FRET-based sensors (24–26) may not provide an accurate description of the crowding experienced by large complexes, such as ribosomes, in the intracellular environment. Yet, given the importance of such large complexes for cellular activity, it is relevant to explore how crowding constrains their mobility *in vivo*.

In this study, we systematically estimated the contribution of three classes of cellular components (mRNA, DNA, and remaining components, i.e., mostly proteins and ribosomes) to the crowding experienced by an encapsulin-based exogenous particle in the cytoplasm of *E. coli* . The particle used has a diameter of 40 nm, approximately the size of a ribosome dimer (27). By measuring the translational diffusion coefficient of this particle after removal of specific macromolecules by either drug treatment or genetically induced depletion, we could estimate the relative contribution of each of the three cellular components to the crowding experienced by the said particle. We observed that the contribution of all three classes is comparable in magnitude but is largest in the case of proteins and ribosomes. Our findings also reveal that, despite their much lower volumetric abundance in the cell, nucleic acids have a non-negligible impact on crowding. We further observed that crowding due to any of the three classes of cellular components shows only a slight dependency on growth conditions. This study shows how different cellular components contribute to crowding in the cytoplasm and, specifically, how they impact the mobility of large macromolecular complexes *in vivo*.

## Materials and Methods

### Plasmids

To express the self-assembling, encapsulin-based 40 nm fluorescent particle derived from the archaeon *Pyrococcus furiosus,* PfVS (17), in *E. coli* , we constructed the pTet-PfVS plasmid, which expresses the particle from an anhydrotetracycline-inducible promoter. This plasmid was constructed by Gibson Assembly (28) after amplification of the *PfVS* gene from pRS306-URA3-PHIS3-PfV-GS-Sapphire (17) by high-fidelity PCR with primers designed with 5’ overhangs complementary to the plasmid backbone (5’-gaaaagaattcaaaagatctATGCTCTCAATAAATCCAAC-3’ and 5’-cctggagatccttactcgagTTATTTGTACAATTCATCAATACC-3’).

The pSN1 plasmid (pBAD24-ISce-1), which encodes the ISce-1 restriction enzyme under the regulation of an arabinose-inducible promoter (29), was obtained as a kind gift from Dr. Christian Lesterlin (CNRS-INSERM; Lyon, France).

### Strains

Throughout this study, the *E. coli* strain LY177 was used. The strain is derived from MG1655 by deletion of *recA* and introduction of two ISce-1 restriction sites, at both *ori* and *ter* chromosomal loci (Δ*recA-Tc ydeO::I-Sce1^CS^, ilvA::I-Sce1 ^CS^*; ref. 29). Where noted, the strain BW25113 (30) was used in complementary experiments. For single particle tracking, both strains were transformed with pTet-PfVS.

For measurements in which the removal of chromosomal DNA was desired, cells were additionally transformed with pSN1. Due to a reported genetic instability that results from leaky expression of ISce-1 (C. Lesterlin, personal communication, 2022) the transformation was performed before every set of experiments.

### Preculturing

Cells were streaked on LB-agar plates (supplemented with 2 g/L glucose, in the case of the pSN1-carrying strain), and a single colony was used to inoculate a loosely-capped tube with 2 mL of M9 medium (42.2 mM Na_2_HPO_4_, 22 mM KH_2_PO_4_, 8.6 mM NaCl, 11.3 mM (NH_4_)_2_SO_4_, 1 mM MgSO_4_, 6.3 µM ZnSO_4_, 7 µM CuCl_2_, 7.1 µM MnSO_4_, 7.6 µM CoCl_2_, 100 µM CaCl_2_, 60 µM FeCl_3_, 2.8 µM thiamine hydrochloride) supplemented with either (i) 5 g/L glucose, (ii) 5 g/L glucose + 100 µg/mL L-arginine, or (iii) 5 g/L glucose + 2 g/L casamino acids (Formedium). The cultures were incubated overnight at 37 °C, with orbital shaking at 300 rpm, after which they were diluted into flasks (100 mL nominal volume) filled with 10 mL of the appropriate culture medium, preheated to 37 °C. Approximately eight hours later, and while still being in exponential phase, cells were diluted once more, but this time split into two separate flasks, each containing 10 mL of culture medium. These dilutions were adjusted based on the known growth rates and incubation period, so that cells remained at a concentration below 3 x 10^8^ cells/mL. A different chemical was added to each flask, as described below. Cell counts were monitored by flow cytometry (BD Accuri C6) and sampling for microscopy imaging was done at a concentration of ∼1.5 – 3 x 10^8^ cells/mL.

For single particle tracking experiments, no inducer (anhydrotetracycline) was added to the cultures since leaky expression from the pTet-PfVS plasmid was sufficient for visualization of the fluorescent particles.

### In vivo perturbation of crowder levels

To measure diffusion under various levels of intracellular crowding, cells grown under each of the three medium conditions (minimal medium supplemented with glucose, glucose and L-arginine or glucose and casamino acids) were subjected to three sets of treatments. To ensure the effect on diffusion was caused by the addition of a given chemical (DMSO, rifampicin or L-arabinose), the cultures were split in two, with one of the flasks being used a negative control.

To probe diffusion in the presence of all cellular crowders, the LY177+pTetPfVS strain was used. 100 µL of DMSO (100%) were added to one flask. As a negative control, no chemical was added to the second flask. Samples were taken ∼45 min after addition of the chemical.

To probe diffusion in the absence of mRNA, the LY177+pTetPfVS strain was used. 100 µL of rifampicin (50 mg/mL; dissolved in DMSO) were added to one flask. As a negative control, the same volume of DMSO was added to the second flask. Samples were taken ∼45 min after addition of the chemicals.

To probe diffusion after DNA depletion, the strain LY177+pTetPfVS+pSN1 was used. 100 µL of DMSO and 180 µL of L-arabinose (10% (m/v)) were simultaneously added to one flask. For comparison, expression of ISce-1 was transcriptionally blocked by simultaneously adding 100 µL of rifampicin and 180 µL of L-arabinose to the second flask. In a variation of this experiment, DMSO or rifampicin were added one hour after L-arabinose. To enable complete degradation of the chromosome (29), samples were taken two hours after addition of L-arabinose.

### Microscopy imaging

After the incubation periods described above, cells were harvested and concentrated 5 – 10-fold by centrifugation (11’000x g, 30 s) of 700 µL of the suspension in 0.22 µm centrifuge tube filters (Costar), followed by resuspension in a fraction of the flow-through. Concentrated cell suspensions were then spotted on glass coverslips (#1.5 thickness, 0.16 – 0.19 mm; Epredia), and the droplets were rapidly covered with agarose pads (preheated to 37 °C). These pads were made of M9 medium and 1.5% (m/v) agarose, supplemented with the same carbon source, and at the same concentration, used in the preculturing (see “Preculturing” above). The appropriate carbon sources were added to a mixture of M9 and agarose previously molten in a microwave oven.

Imaging was done using an Eclipse TiE inverted microscope (Nikon Instruments) equipped with a 100x Plan Apo Lambda NA 1.45 objective (Nikon Instruments), with additional 1.5x optical magnification. The system was coupled to an iXon Ultra 897 EMCCD camera (Andor) and an LED light source (Aura Light II, Lumencor). Two filter sets were used, both containing a 495 nm dichroic mirror and a 525/50 nm emission filter, and either a 470/40 nm or 390/40 nm excitation filter (AHF Analysentechnik). First, snapshots were taken with the brightfield channel for cell segmentation. Cells were then exposed to 395 nm light (LED intensity: 100%) for 2 s, as this was found to have a photoactivation effect, increasing the brightness of the PfVS particles when later excited with 485 nm light. Finally, image acquisition was done under excitation at 485 nm (LED intensity: 100%), for a total of 100 frames, with an exposure time of 19.4 ms per frame, readout at 17 MHz and EM gain of 100.

### Single particle tracking

Single particle tracking and determination of the mean squared displacement, *MSD*, of each particle were done using the functions *locate()* and *msd()* of the *trackpy* package (31), respectively, with the following settings: *gap=5 px, diameter=9 px, mem=1 frame, minlength=33 frame, noise_size=1, dilate=5* . An automatic filtering step was applied to remove spurious trajectories based on the properties reported by the tracking routine, namely the shape (mean *eccentricity* < 0.2), size (mean *size* < 2.1) and precision of localization (median *ep* < 0.31) of the detected particles.

Cells and corresponding particle trajectories were rotated to a common orientation (long cell-axis parallel to the x-axis, and short cell-axis parallel to the y-axis) using the *ColiCoords* package (32). Thereafter, the one-dimensional mean squared displacement, *MSD_x_*, was obtained using the longitudinal component of the particle trajectories. Diffusion coefficients were obtained for every tracked particle by fitting *MSD_x_* to the equation:

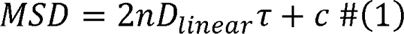

where *D_linear_* is the linear diffusion coefficient; *n* is the number of dimensions (*n* = 1, in the case of one-dimensional diffusion); c is an offset used to account for errors in the localization (33).

### Estimation of diffusion coefficients in crowder-less environments

The unhindered diffusion coefficient, *D_o_*, of particles moving in a water was estimated using the Stokes-Einstein equation for spherical objects (34),

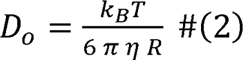

where *k_B_* is the Boltzmann constant (1.38 x 10^-23^ J.K^-1^), *T* is the absolute temperature (310.15 K), *R* is the radius of the diffusing particle (40 nm/2 = 20 nm), and *η* is the viscosity of the fluid (6.9 x 10^-4^ Pa s, the viscosity of water at 37 °C; ref. 35).

### Correction of diffusion coefficients for the effect of confinement

To correct the experimentally determined diffusion coefficients (obtained as described in “Single particle tracking”) for the effect of confinement, an iterative computational approach was implemented (Fig. 2C). In each iteration of this method, the particle-based simulator software *Smoldyn* (36) was used to generate random trajectories in spherocylindrical compartments.

First, the dimensions of the spherocylindrical compartments were chosen to match the morphology of *E. coli* cells under the relevant growth condition. Specifically, for the correction of diffusion coefficients obtained in glucose-grown cells, the compartments were defined with a radius and length of 0.45 µm and 2.6 µm, respectively. For growth on glucose and L-arginine, a radius and length of 0.425 µm and 2.8 µm were used; for growth on glucose and casamino acids, the same parameters had values of 0.475 µm and 3.2 µm (Fig. S1).

In each iteration, trajectories were generated using parameters that closely matched our experimental imaging setup. In particular, 300 trajectories were simulated, each containing 100 localizations, with a time step of 19.4 ms. To generate these trajectories, the value of the real diffusion coefficient, i.e., unaffected by confinement, *D_o_*, had to be provided as input. Since *D_o_* is *a priori* unknown, a value of 1.2 x 10^log10(*Dexp)*^, where log_10_(*D_exp_*) is the experimentally determined value, was used as the initial guess.

For each simulated trajectory, the one-dimensional *MSD* was obtained based on the longitudinal coordinate (x), and the corresponding diffusion coefficient, *D_pred_*, was obtained using Eq. Error! Reference source not found.. The population average of log_10_(*D_pred_*) values was then compared with the experimentally-determined log_10_(*D_exp_*) values. The squared error between the latter two values, ε^2^,

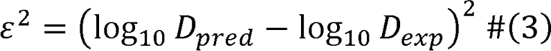

was then iteratively minimized by adjusting the value of the input parameter *D_o_*. This optimization was implemented with the *scipy.optimize.minimize()* function, using the Nelder-Mead solver, and with parameters *fatol=1e-4* and *maxfev=50* (37).

## Results

### Overview of the experimental approach

To quantify how various macromolecules hinder the mobility of a 40 nm fluorescent particle (17), we sought to determine the diffusion coefficient of this particle under different conditions of intracellular crowding, ranging from complete absence of crowders to presence of all cellular crowders. Specifically, using the Stokes-Einstein equation, we first estimated the diffusion coefficient in a scenario where macromolecular crowding is absent (Fig. 1, bottom). Second, we performed *in vivo* single particle tracking to measure the diffusion coefficient of the same particle under unperturbed conditions, when all crowders are present (Fig. 1, top). Finally, we determined the diffusion coefficient after selectively removing mRNA, by rifampicin treatment, and DNA, by genetically induced DNA degradation (Fig. 1, middle). Calculating the differences between the diffusion coefficients across these various investigated scenarios, we obtained the effective contribution of each class of cellular components to the crowding experienced by the 40 nm particle used (Fig. 1, right).

**Figure 1.**
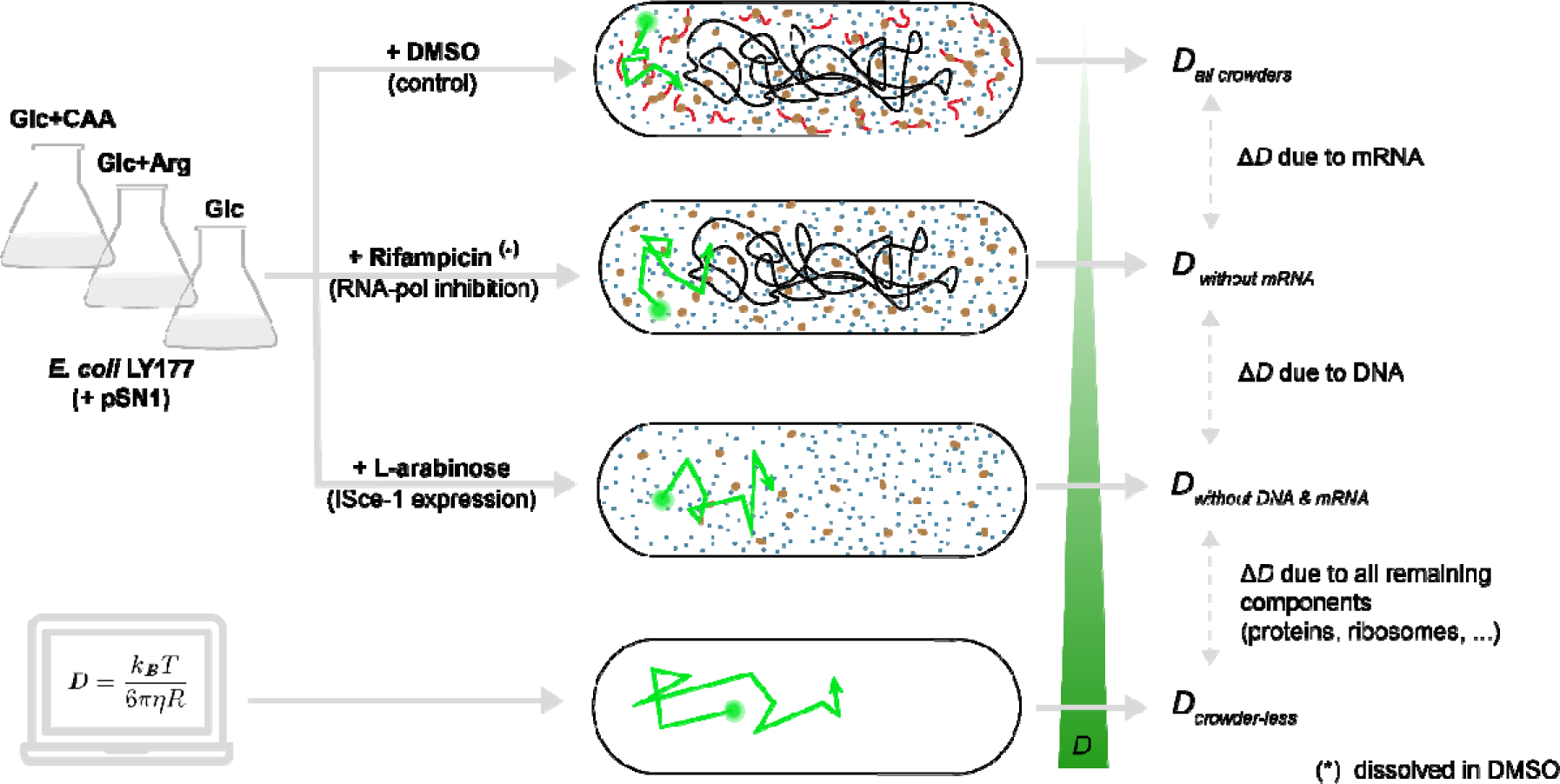
Overview of the experimental approach. *E. coli* LY177 cells expressing the 40 nm fluorescent particle (PfVS) were cultivated in minimal medium supplemented with glucose (Glc), L-arginine (Arg) and/or casamino acids (CAA) and were imaged under the microscope for single particle tracking. DMSO was added as a control treatment for measurement of the diffusion coefficient in cells containing all macromolecular components, *D_all crowders_* (top). The treatment with rifampicin (dissolved in DMSO) was used to block RNA-polymerase activity and to thereby allow measurement of the diffusion coefficient in cells depleted of mRNA*, D _without mRNA_* (second from top). L-arabinose was added to cells additionally carrying the pSN1 plasmid to induce expression of ISce-1; this treatment allowed measurement of the diffusion coefficient in cells depleted of both DNA and mRNA, *D_without DNA & mRNA_* (third from top). To estimate the diffusion coefficient in the absence of macromolecular crowders, *D_crowder-less_,* the Stokes-Einstein equation was used, assuming that the solvent is water at 37 °C (bottom). PfVS particle: green circle; DNA: black lines; mRNA: red lines; ribosomes: large brown circles; proteins: small blue circles.

### Approach to correct diffusion coefficients for the effect of confinement

For a 40 nm particle diffusing in water at 37 °C, the Stokes-Einstein equation (Eq. 2 in Materials and Methods) predicts a diffusion coefficient, D, of 16.4 µm /s. A particle diffusing with such a value of D would, in a period of 100 ms, which corresponds to the timescale used in our in vivo single particle tracking measurements (see below), cover a distance of approximately 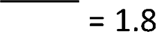 µm in one dimension. The similarity between the latter value and the size of an E. coli cell (Fig. S1) indicates that confinement by the cell membrane could lead to an underestimation of the diffusion coefficients obtained by single particle tracking.

To confirm whether confinement could affect our measured diffusion coefficients, we ran computational simulations with Smoldyn (36) to obtain random trajectories of the particle in cell-shaped compartments (Fig. 2A). Specifically, we simulated 300 trajectories with the diffusion coefficient obtained from the Stokes-Einstein equation for a 40 nm particle moving in water (16.4 µm^2^/s). We then determined the mean squared displacement (MSD) for each of these trajectories and, assuming a linear dependence of *MSD* on time (Eq. 1 in Materials and Methods), estimated the *apparent* diffusion coefficients. We found that these diffusion coefficients indeed decreased with decreasing “cell” lengths (Fig. 2B). This finding confirms that, by determining diffusion coefficients from observed trajectories, we could potentially underestimate the *true* diffusion coefficient – i.e., the value obtained in an otherwise unconfined environment.

**Figure 2.**
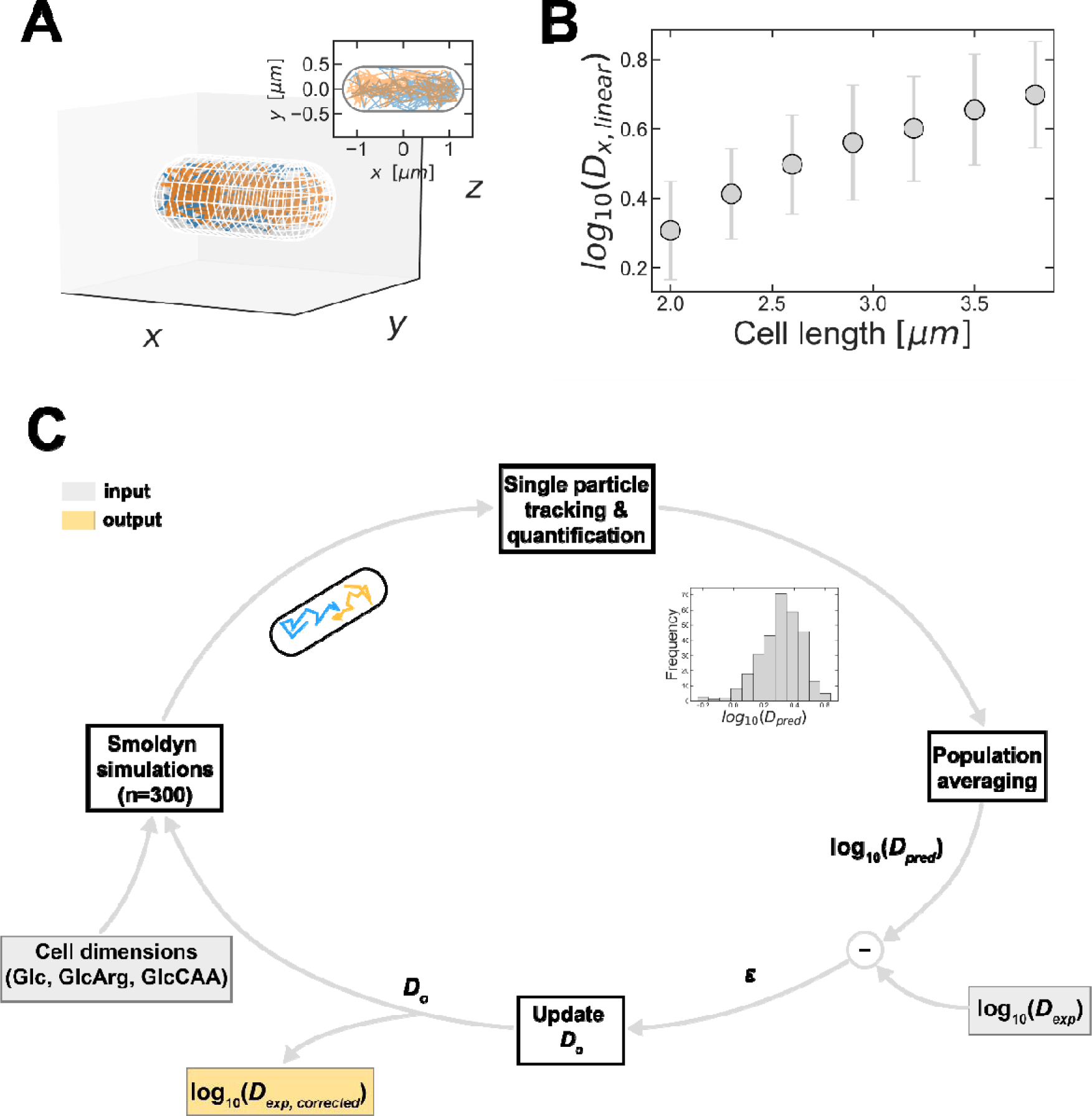
Underestimation of diffusion coefficients due to confinement by the cell membrane is corrected with a computational procedure. (**A**) Visualization of two examples of simulated 3D-particle trajectories (blue and orange lines) in a spherocylindrical compartment with dimensions comparable to an *E. coli* cell grown on glucose (length: 2.6 µm; radius: 0.45 µm). The unconfined diffusion coefficient, *D_o_* = 16.4 µm^2^ /s, was obtained using the Stokes-Einstein equation (Eq. **Error! Reference source not found.** ), with *T* = 310.15 K, *R* = 20 nm and *η* = 6.9 x 10^-4^ Pa.s. Inset: projection of the two trajectories on the x-y plane. (**B**) Longitudinal diffusion coefficients obtained from simulated particle trajectories in spherocylindrical compartments (“cells”) with a radius of 0.45 µm and variable length. The unconfined diffusion coefficient was set to *D_o_ =* 16.4 µm^2^/s. Each marker shows the mean ± standard deviation of n = 300 trajectories. (**C**) Iterative procedure to determine the value of *D_o_* (diffusion coefficient in the absence of confinement) from an experimentally determined value, *D_exp_*. In each iteration, 300 particle trajectories are simulated in a spherocylindrical compartment with dimensions matching those of cells grown with the appropriate carbon source (Glc, GlcArg or GlcCAA growth conditions). For each trajectory, the diffusion coefficient is obtained by fitting the corresponding mean squared displacement to Eq. **Error! Reference source not found.** . The population-averaged log_10_*D* value of the simulated trajectories, log_10_(*D_pred_*), is compared with the experimental value, log_10_*D_exp_*. If the residual, *ε*, exceeds a specified threshold value, the value of *D_o_* is updated; otherwise, *D_o_* is accepted as the counterpart value of *D_exp_* corrected for confinement.

We then set out to eliminate the effect of confinement from the diffusion coefficients we obtained from *in vivo* measurements. We therefore implemented an iterative computational methodology to estimate the diffusion coefficients in unconfined environments from the values obtained under confinement (Fig. 2C) – an approach which is similar to one that has been recently described (38). Briefly, we started with a first guess of the unconfined diffusion coefficient, *D_o_*. With this value, we simulated particle trajectories in a compartment with the same shape as *E. coli*. The dimensions of these “cells” were chosen to match the morphology observed experimentally under each growth condition (Fig. S1). From the *MSD*s of these simulated trajectories, we estimated the corresponding *apparent* diffusion coefficient. The value of *D_o_* was then automatically adjusted by an optimization algorithm until the population-averaged apparent diffusion coefficient matched the experimentally determined values. At this point, *D_o_* was taken as the corrected counterpart of the experimental values, i.e., corrected for the effect of confinement.

### Determination of diffusion coefficients in crowded environments

Having established a way to correct the diffusion coefficients obtained from experimental measurements for the effect of confinement, we then proceeded to measure diffusion *in vivo* in the presence of all macromolecular crowders. For this, *E. coli* was grown in minimal medium with either glucose (Glc), glucose and L-arginine (GlcArg), or glucose and casamino acids (GlcCAA). Cells from these cultures were sampled while in the exponential growth phase, were placed on microscopy slides and were covered with agarose pads made of the same medium as the liquid cultures. With fluorescence microscopy, we imaged the cells at a rate of 51.6 frames per second. We then tracked the fluorescent particles in the microscopy movies with a tracking algorithm (31) and determined the confinement-corrected diffusion coefficients. In most cases, however, the difference between the experimental- and corrected values was minimal (Fig. S2).

Following this procedure, we obtained diffusion coefficients that were more than two orders of magnitude lower than the values predicted by the Stokes-Einstein equation for a crowder-less environment (Fig. 3A), which is consistent with the cytoplasm being a heavily crowded environment. We found that the diffusion coefficients varied slightly across growth conditions, being lowest on glucose and highest on glucose with casamino acids, which are two conditions with an almost two-fold difference of growth rate (0.48 h^-1^ and 0.95 h^-1^, respectively; Fig. 3A). We further observed that different *E. coli* strains showed different diffusion coefficients under the same growth conditions. With the strain LY177, we obtained higher values than with the strain BW25113, while both strains displayed the same condition-dependency (Fig. S3). These results show that crowding by all macromolecules in a cell reduces the mobility of the 40 nm particle by more than two orders of magnitude.

**Figure 3.**
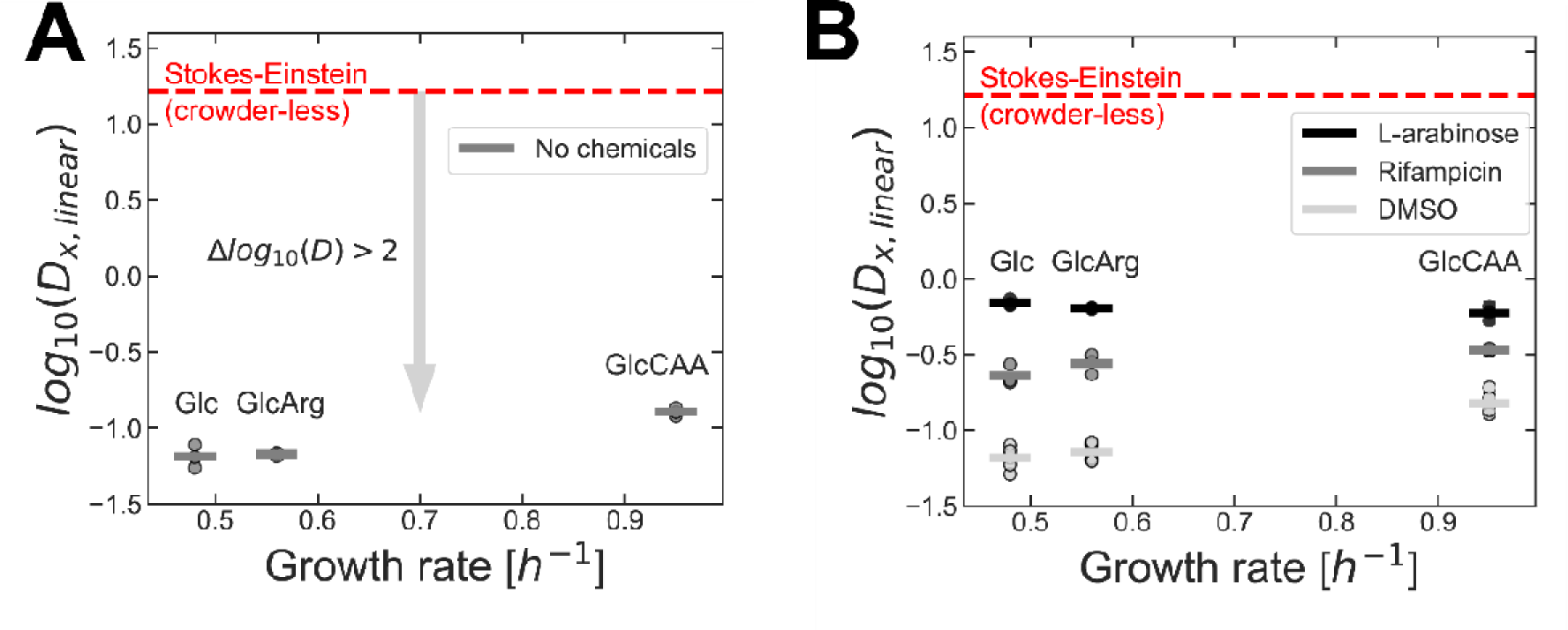
Removal of mRNA and/or DNA leads to an increase in the diffusion coefficient. (**A**) Diffusion coefficients measured in cells grown in different media (Glc, GlcArg, GlcCAA), corrected for the effect of confinement, as shown in Fig. **Error! Reference source not found.** C. The measurements were done in unperturbed cells (i.e., not exposed to any chemicals). (**B**) Same as (**A**), but the diffusion measurements were done after various treatments (“DMSO”, “Rifampicin” and “L-arabinose”). With the exposure to DMSO, cells retained all macromolecular crowders and the diffusion coefficients reproduced the values obtained in unperturbed cells (note the similarity with (**A**); see also comparison of the values in Fig. **Error! Reference source not found.** A and B). With the exposure to rifampicin (dissolved in DMSO), mRNA was degraded and the diffusion coefficient increased relative to the control condition (DMSO). Addition of L-arabinose to LY177+pSN1 led to the induction of ISce-1 expression, resulting in the degradation of chromosomal DNA and in a further increase in the diffusion coefficients. In both (**A**) and (**B**), averages of values determined in n ≥ 3 experiments (markers) are shown by the horizontal bars. The horizontal red line displays the predicted diffusion coefficient of the 40 nm particle in water (i.e., crowder-less environment), calculated with the Stokes-Einstein equation (Eq. 2).

To assess how much mRNA contributes to the crowding experienced by the particle, we repeated the measurements after treating cells with rifampicin, dissolved in DMSO. By binding to, and blocking the activity of, RNA-polymerase, rifampicin halts all transcriptional activity in the cell (39). Given the short lifetime of mRNA (40–42), rifampicin treatment leads to the depletion of the cellular pool of that macromolecule (43, 44). After depleting mRNA with this treatment, we observed that the diffusion coefficient of the particle increased significantly, compared to corresponding negative controls, i.e., exposure to DMSO, under all three growth conditions (Fig. 3B; “Rifampicin” vs. “DMSO”; *p* ≤ 6.1 x 10^-5^, two-sided t-test for independent samples). Importantly, the negative controls were themselves comparable to untreated conditions (Fig. S4A and B). These results reveal that despite its low abundance, accounting for only 0.8% of the cell dry weight (as estimated in Supplemental Note 1.3), mRNA is responsible for a non-negligible part of the hindrance experienced by the 40 nm particle in vivo.

Next, we focused on assessing the contribution of DNA to crowding. For this, we used a strain carrying the arabinose-inducible pSN1 plasmid. This plasmid encodes a restriction enzyme, ISce-1, that targets two restriction sites in the chromosome of LY177, where it introduces DNA double-strand breaks. Given the absence of the DNA repair enzyme RecA in this strain, action of endonucleases then leads to degradation of the entire chromosome (29).

We first addressed concerns that leaky expression of the ISce-1 enzyme could be detrimental for cell physiology (C. Lesterlin, personal communication, 2022) by confirming that cells with and without the pSN1 plasmid grew with indistinguishable growth rates (Fig. S5). Then, to confirm that the expression of ISce-1 leads to complete degradation of chromosomal DNA, we stained cells with the fluorescent DNA-binding dye DAPI. Two hours after addition of the inducer, L-arabinose, almost all cells were non-fluorescent, in contrast to a control condition in which the expression of ISce-1 was transcriptionally blocked (Fig. S6A, top row; Supplemental Note 2).

After confirming that growth rate was unaltered by the presence of pSN1, and that DNA could be successfully degraded upon expression of ISce-1, we performed single particle tracking in cells expressing this enzyme, two hours after induction with L-arabinose. Consistent with our expectation that the removal of DNA should lead to an increase in diffusion, the diffusion coefficients observed with this treatment were even higher (*p* ≤ 8.9 x 10^-4^, two-sided t-test for independent samples) than in mRNA-depleted cells (Fig. 3B; “L-arabinose” vs. “Rifampicin”).

We hypothesized that cells with degraded DNA were also free of mRNA. First, lack of DNA means that no template molecule exists from which mRNA transcripts can be synthesized. Second, mRNA has a lifetime on the order of a few minutes (40–42), so that any molecules formed prior to the degradation of DNA are likely degraded in the course of a typical treatment (two hours). To test this hypothesis, we performed experiments in which either rifampicin or DMSO were added to cells *already* expressing ISce-1. With this approach, we sought to determine the impact of enforced mRNA depletion (by rifampicin) in cells without DNA. We added either chemical one hour after induction of ISce-1 expression with L-arabinose, and confirmed that DNA was absent in both cases (Fig. S6A, bottom row). Here, the treatment with rifampicin did not result in increased diffusion coefficients, when compared to the control treatment with DMSO (Fig. S4B, bottom). In line with our hypothesis, this suggests that DNA degradation by ISce-1 is sufficient to cause the disappearance of cellular mRNA. Considering this finding, the difference between the diffusion coefficients measured under the two treatments described earlier (Fig. 3B; “L-arabinose” vs. “Rifampicin”) is due to the presence/absence of DNA (see Fig. 1).

Finally, we compared the diffusion coefficients obtained after ISce-1 expression with the value estimated using the Stokes-Einstein equation (Fig. 3B; “L-arabinose” vs. “Stokes-Einstein”). Notably, after the combined removal of DNA and mRNA that results from the expression of ISce-1, the measured diffusion coefficients were still much lower than the value predicted by the Stokes-Einstein equation for an aqueous solution. This difference indicates that other cellular components, namely proteins and ribosomes, strongly hinder particle motion. It is, however, unclear whether this hampering effect results from a high viscosity caused by small molecules (compared to the 40 nm particle) or from the presence of obstacles of size comparable to the size of the particle.

### Contribution of various cellular components to crowding

With the diffusion coefficients determined after depletion of the various macromolecular crowders (Fig. 3B), we could then estimate the relative contribution of each of the latter to the crowding experienced by the 40 nm particle. To achieve this, we started by determining the difference between log_10_*D* values of conditions that differed by presence/absence of any one of three classes of cellular components (i.e., DNA, mRNA and remaining cellular components). Here, the contribution of mRNA was estimated from the difference between the rifampicin-treatment and the DMSO control; the contribution of DNA from the difference between the ISce-1-induction and the rifampicin-treatment; and the contribution of the remaining cellular components from the difference between the crowder-less and ISce-1-induction conditions (Fig. 1). Then, these values were normalized by the contribution of all cellular crowders, defined as the difference between the two “extreme” scenarios (i.e., crowder-less condition and DMSO control).

Following this approach, we found that the major contribution to crowding stemmed from cellular components other than DNA and mRNA, accounting for ∼63% of the reduction in the diffusion coefficient of the particle (average across the Glc, GlcArg and GlcCAA growth conditions; Fig. 4A). Furthermore, while the exact values differed slightly across the growth conditions tested, the contributions of mRNA and DNA were estimated at ∼22% and ∼16%, respectively (averages across the Glc, GlcArg and GlcCAA growth conditions; Fig. 4A). In particular, the contribution of DNA exhibited a monotonic decreasing trend with growth rate, whereas the contribution of the “other” cellular components followed a linear increasing trend with growth rate (Fig. 4A). Taken together, these results reveal that all three classes of cellular components considered here contribute to the crowding experienced by the 40 nm particle with comparable magnitude, each accounting for >16% of the reduction in diffusion, and with slight condition-dependency.

**Figure 4.**
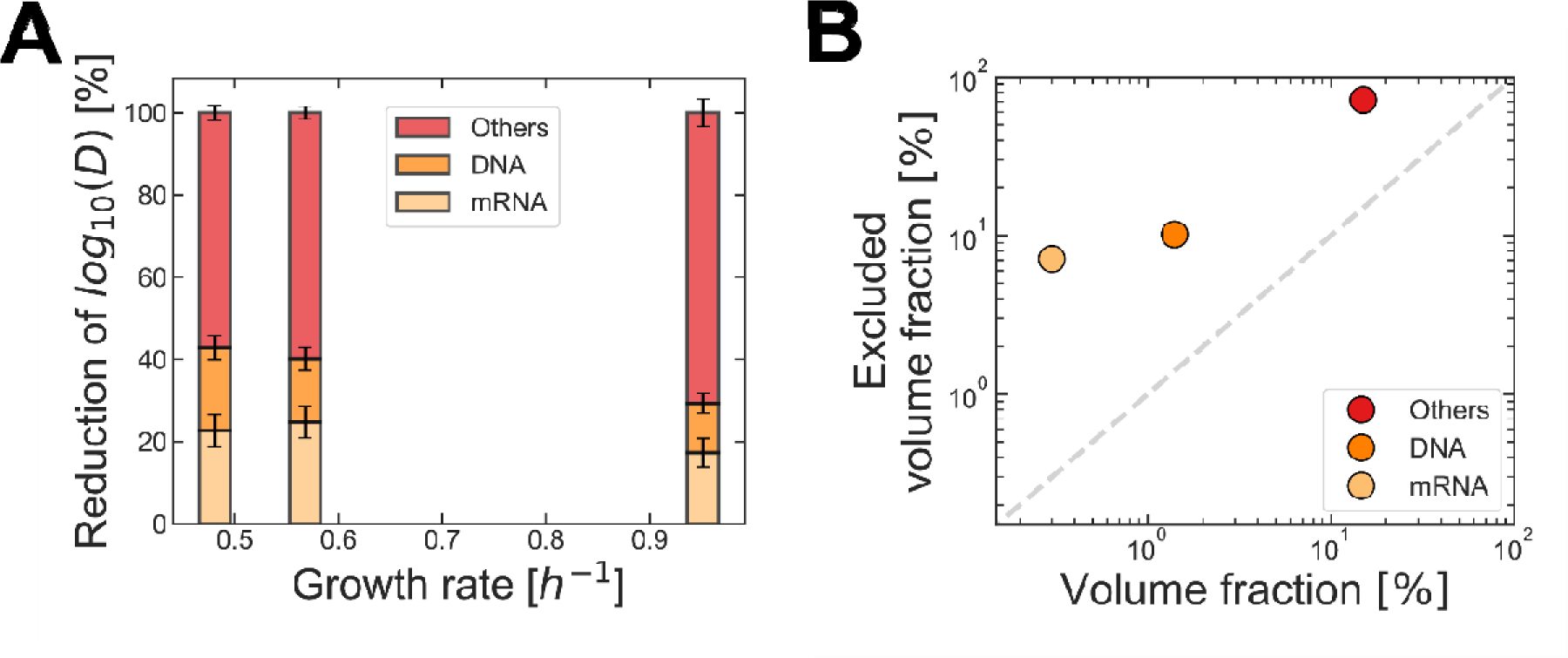
Different cellular components have a comparable effect on crowding experienced by the particle despite their widely different volumetric fractions. (**A**) Relative contribution of mRNA, DNA and remaining cellular components to the decrease of diffusion coefficient of the 40 nm particle. The values were calculated as differences between confinement-corrected log_10_*D* values: for mRNA, between the rifampicin- and DMSO-treatments; for DNA, between the L-arabinose- and rifampicin-treatments; for the remaining cellular components, between the Stokes-Einstein diffusion coefficient and the values obtained after L-arabinose treatment (see Fig. 1). All values were normalized by the difference between log_10_*D* in the two extreme scenarios in terms of crowding (i.e., Stokes-Einstein and DMSO-treatment). The bars were placed on the x-axis according to the average growth rate under the different growth conditions (Glc, GlcArg, GlcCAA) prior to any of the treatments described. Error bars were obtained by propagation of the uncertainty associated with each value of log_10_*D*. (**B**) Estimated fraction of cellular volume excluded to the 40 nm particle plotted against the estimated actual volume fraction of a given cellular component. The y-values were obtained from the differences between log_10_*D* values under the various experimental treatments (growth condition: Glc), and through application of Eq. S2.4 (Supplemental Note 3). Additivity of excluded volumes due to the various cellular components was assumed (Supplemental Note 3). The corresponding error bars (not visible) are the analytically-derived uncertainties. The x-values are reported (22) or calculated (Supplemental Note 1) values of the volume fractions of DNA, mRNA and proteins+ribosomes. Unlike the excluded volumes, these values do not depend on the particle used as diffusion probe, but rather on the molecular composition of the cell.

To translate our experimental diffusion measurements into apparent volumes taken up by each of the cellular component (mRNA, DNA and others), we made use of a model relating diffusion coefficients to the fraction of excluded volume caused by crowders in suspension (45). In a simplified description of crowding in the cell, we assumed that excluded volumes are additive, and that crowding is the only cause of the reduction in the diffusion coefficient (Supplemental Note 3). These model predictions show that, for cells grown on glucose, the excluded volume caused by either DNA or mRNA corresponds to ∼10% and ∼7% of the entire cell volume, respectively (Fig. 4B). Both values are larger than the volume fractions of DNA or mRNA in the cell (1.4% and 0.3%, respectively), which can be estimated from the known abundance of these molecules and their specific volumes (ref. 22 and Supplemental Note 1.1). Overall, more than 80% of the cellular volume seems to be inaccessible to the particle under each of the growth conditions studied (Fig. 4B). This value is significantly larger than previous estimates of the total fraction of cellular volume occupied by macromolecules, ranging from 13% to 24% (4, 24). That large excluded volume also reveals the mobility constraints imposed on particles/complexes of approximately 40 nm in the cytoplasm of E. coli.

## Discussion

By measuring the diffusion coefficient of a 40 nm particle in the cytoplasm of E. coli upon depletion of DNA and/or mRNA and estimating it in a crowder-less environment, we determined the effective crowding contribution by DNA, mRNA and the remaining cellular components. Furthermore, by applying a simple model that describes diffusion in crowded environments, we estimated the fraction of cellular volume that is inaccessible to the cytoplasmic particle due to each of the three classes of cellular components considered. We found that crowding resulting from each of these classes of components is comparable in magnitude despite the large differences in the actual volume fraction of these crowders in the cytoplasm. The values display a slight degree of condition-dependency but, in all cases, crowding is dominated by the effect of components other than mRNA/DNA, such as proteins and ribosomes.

The fact that both DNA and mRNA exert a measurable hindering effect on the mobility of the particle is surprising given that, individually, each of these nucleic acids accounts for less than 2% of the cell volume (ref. 22 and Supplemental Note 1.1). Indeed, with the assumption that these molecules only hinder the diffusion of the particle due to excluded volume effects, our model-based estimates indicate that both DNA and mRNA render at least 7% of the cellular volume inaccessible to the particle (Fig. 4B). The discrepancy between these two values stresses that *real* and *excluded* crowder volumes may differ (4, 6). Particularly for linear macromolecules such as DNA and mRNA, with a diameter much smaller than the particle, a fully expanded conformation is expected to pose less hindrance to the motion of the particle than a more folded (“mesh-like”) conformation. Thus, the condensation state of these molecules, i.e., the degree to which they are spatially compacted, could potentially account for part of the difference between the real and excluded volumes.

In the case of DNA, the disproportionally large excluded volume, compared to the “real” volume occupied by this molecule, is easily understood when one considers that DNA is the core constituent of the nucleoid (46). The latter is a condensed polymeric structure with a mesh size of approximately 50 nm (15). Since this number (50 nm) is merely descriptive of the average nucleoid structure, it is likely that local compaction of the “mesh” creates regions inaccessible to the 40 nm particle used in this study. Such local compaction may thus be one of the main reasons why the excluded volume of DNA exceeds the real volume of the nucleoid. Similarly, condition-dependent changes in the volume excluded by DNA may arise from more global, as opposed to local, changes in the nucleoid structure, namely from changes in the average mesh size. In this regard, evidence suggesting that nucleoid compaction decreases with increasing cell size (14) may play a role in explaining why the estimated excluded volume of DNA under the condition of largest cell dimensions (glucose with casamino acids; Fig. S1) was the smallest among the three experimental conditions (Fig. 4A).

In turn, mRNA molecules are much shorter (typically 750 nucleotides long; ref. 47) than DNA, so that considerations about a “mesh-like” polymeric structure may not be valid. How can we then explain the seemingly large excluded volume caused by these low abundant macromolecules? To answer this question, two aspects must be considered. First, recent evidence shows that treatment with rifampicin may lead to the removal not only of mRNA but also of a fraction of rRNA, particularly of the 16S and 23S types (refs. 48, 49; see also Fig. S7). Consequently, the observed increase in diffusion coefficient (Fig. 3B) may partly result from degradation of ribosomes. Secondly, interactions of mRNA with other cellular components may also play a role in explaining the apparently large excluded volume of mRNA. For example, by undergoing association with multiple ribosomes, mRNA and ribosomes can together form large structures, named polyribosomes (50). Another example is the involvement of mRNAs in the “transertion mechanism”, in which simultaneous transcription, translation and insertion of proteins in the plasma membrane contributes to the radial expansion of the nucleoid (12, 51). The interference of rifampicin with both of these interactions, i.e., disruption of “transertion” and disassembly of polyribosomes, could be reasons behind the observed expansion of the nucleoid upon exposure to this chemical (11, 12, 19, 52). Irrespective of the true mechanistic explanation for the observed nucleoid expansion, this behavior exemplifies how experimental depletion of cellular mRNA not only removes one potential crowder from the cell, but also how it can lead to a more widespread reorganization of the cytoplasm. In this study, we estimated the excluded volume fraction of mRNA based on an observed increase in the diffusion coefficient of the particle upon removal of this molecule. Yet, the diffusion coefficient is a readout for all the changes taking place in the cytoplasm. Thus, the excluded volume estimate should be seen as an aggregate measure that also includes the effects resulting from the interaction of mRNA with other components, as described above. As such, it is not entirely surprising that the apparent excluded volume caused by mRNA largely exceeds the real volume of this macromolecule. Still, the excluded volume fraction we estimated provides a more realistic depiction of the effect that mRNA, directly and indirectly, has on the mobility of the particle.

Our experiments suggest that the contribution of the remaining cellular components, such as proteins and ribosomes, to the crowding experienced by the 40 nm particle is the most prevalent among the classes studied (Fig. 4A). This observation qualitatively aligns with the results of a previous study which showed that ribosomes are the major cytoplasmic crowders affecting the mobility of the same particle (PfVS) in yeast (17). In *E. coli* , however, the lack of DNA compartmentalization, together with the large size of the particle used (40 nm) had led us to expect a lower *relative* contribution of ribosomes/proteins, at least compared to that of DNA. These results were therefore somewhat unexpected. They may nevertheless be explained by the high abundance of proteins and ribosomes, which jointly amount to more than 70% of the cell dry weight (or ∼15% of the cell volume; Supplemental Note 1.2 and 1.4). The high abundance of both small-(individual proteins) and large obstacles (ribosomes and, potentially, protein agglomerates) is clearly more effective at constraining the mobility of 40 nm particles than the large nucleoid at the center of the cell.

Had the mean squared displacements obtained in this work been fit with a power-law equation (i.e., *MSD* ∼ *τ^α^*) instead of assuming a linear dependence on time (*MSD* ∼ *τ^α^*, as in Eq. 1), we would have obtained values of *α* < 1 under the condition of highest crowding (DMSO control; Fig. S8). Such values can indicate that particles undergo subdiffusive motion in this condition. Yet, the fact that *α* varies across conditions of differing crowder levels implies that the diffusive regime varies accordingly. These changes in diffusive regime add a layer of complexity to the data, which we dealt with by determining *effective* diffusion coefficients (using Eq. 1). Remarkably, the fact that *α* ≈ 1 after removal of mRNA by itself suggests that this molecule, more than DNA or any other macromolecule, may be responsible for the viscoelastic properties of the cytoplasm at the 40 nm scale. Assuming the view that *α* < 1 is an indication of crowding (53), the values of *α* ≈ 1 when DNA, proteins and ribosomes are still present could mean that these components exert a hindering effect on particle motion that is driven by a change in bulk viscosity rather than crowding. This view is nevertheless disputed (54, 55), and it is unclear which of the two effects, crowding or viscosity, dominates in those experimental conditions. Yet, even if viscosity has to be factored in the analysis, this would not change the fact that proteins and ribosomes remain the most significant cause for the slowdown of diffusion of the 40 nm particle *in vivo*.

## Conclusion

In this study, we have determined the extent to which the hindrance of mobility experienced by a 40 nm particle in the cytoplasm is due to mRNA, DNA and all other cellular components (including proteins and ribosomes). While all these macromolecules exert an effect of comparable magnitude on the diffusion of this particle, it is noteworthy that the hindrance posed by proteins and ribosomes dominates. Increased viscosity could play a role in this reduction of the diffusion *in vivo*, but the formation of larger-sized protein obstacles is another option to be considered. Overall, our measurements shed light into the constraints that limit the mobility of large macromolecular complexes in the cytoplasm of a bacterial cell.

## Author contributions

J.L. performed all experiments and data analysis, under the supervision of M.H. Both authors conceptualized the study and wrote the manuscript.

## Acknowledgements

The authors thank Dr. Christian Lesterlin for providing the LY177 strain and the pSN1 plasmid; Dmitrii Linnik for helping with the implementation of the *Smoldyn* simulations; Jelle de Boer for performing insightful exploratory experiments with the DNA-degradation system; Bert Poolman, Ed Smith and Yusuke Himeoka for helpful discussions. This research was supported by the “BaSyC – Building a Synthetic Cell” Gravitation grant (024.003.019) of the Netherlands Ministry of Education, Culture and Science (OCW) and the Dutch Research Council (NWO), and of the Dutch Research Agenda (NWA) grant (NWA.1292.19.170) financed by the Dutch Research Council (NWO).

## Supporting citations

References (56–64) appear in the Supporting Material.

