## Supplemental Information for "Contribution of cellular macromolecules to the diffusion of a 40 nm particle in *Escherichia coli*"

### Supplemental Note 1 – Estimation of macromolecule mass and volume fractions

##### 1. Volume fraction of mRNA

The cellular concentration of mRNA, $c_{mRNA}$, is determined by:

$$\begin{aligned} c_{mRNA}=f_{mRNA}\times c_{RNA} \#\left( S1.1 \right) \end{aligned}$$

where $f_{mRNA}$ is the mass fraction of all RNA that is mRNA, and $c_{RNA}$ is the cellular concentration of all RNA. Using the partial specific volume of RNA, $\nu_{RNA}$, we can convert the concentration of mRNA into the corresponding volume fraction, $\phi_{mRNA}$:

$$\begin{aligned} \phi_{mRNA}=\nu_{RNA}\times c_{mRNA}=\nu_{RNA}\times\left( f_{mRNA}\times c_{RNA} \right) \#\left( S1.2 \right) \end{aligned}$$

Using the values from Table S1:

$$\begin{aligned} \phi_{mRNA}=\left( 0.569\frac{mL mRNA}{g mRNA} \right)\times\left( 0.04\frac{mg mRNA}{mg RNA} \right)\times\left( 120\frac{mg RNA}{mL cell} \right)\approx0.3\% \#\left( S1.3 \right) \end{aligned}$$

##### 2. Volume fraction of (all) proteins and ribosomes

On a per-gram dry weight (gDW) basis, the volume occupied by macromolecules of type $i$, is given by:

$$\begin{aligned} V_{i} = \psi_{i}\times\nu_{i} \#\left( S1.4 \right) \end{aligned}$$

where $\psi_{i}$ is the mass fraction of macromolecule $i$ in the dry biomass, and $\nu_{i}$is its partial specific volume.

Similarly, we can obtain the volume occupied by 1 gDW of cellular material, $V_{cell}$, by taking the inverse of the cellular dry density, $\rho_{cell}$:

$$\begin{aligned} V_{cell} = \frac{1}{\rho_{cell}} \#\left( S1.5 \right) \end{aligned}$$

We can then determine the volume fraction of macromolecule $i$in a cell, $\phi_{i}$, as the ratio between the volume occupied by a given macromolecule and the volume of the whole cell:

$$\begin{aligned} \phi_{i} = \frac{V_{i}}{V_{cell}} \#\left( S1.6 \right) \end{aligned}$$

Then, by noting that ribosomes are composed of proteins and rRNA, determining the total volume fraction of proteins and ribosomes in the cell, $\phi_{proteins + ribosomes}$, amounts to simply determining the sum of volume fractions of proteins and rRNA, $\phi_{proteins}$ and $\phi_{rRNA}$, respectively:

$$\begin{aligned} \phi_{proteins + ribosomes} = \phi_{proteins}+ \phi_{rRNA} \#\left( S1.7 \right) \end{aligned}$$

Combining all expressions shown above, and further noting that the rRNA is only a fraction of the total cellular RNA ($\psi_{rRNA} = f_{rRNA}\times\psi_{RNA}$), by using the values from Table S1 we arrive at:

$\begin{aligned} \phi_{proteins+ribosomes} =\frac{\left( \frac{0.86 g rRNA}{g RNA} \right)\times\left( \frac{0.205 g RNA}{g DW} \right)\times\left( \frac{0.569 mL rRNA}{g rRNA} \right) +\left( \frac{0.55 g protein}{g DW} \right)\times\left( \frac{0.728 mL protein}{g protein} \right)}{\frac{1}{0.30 \frac{gDW}{mL cell}}}\approx15\% \#\left( S1.8 \right) \end{aligned}$

##### 3. Mass fraction of mRNA

The mass fraction of mRNA in a cell, $\psi_{mRNA}$, is directly obtained from the mass fraction of all RNA in a cell, $\psi_{RNA}$, and the fraction thereof which is mRNA, $f_{mRNA}$:

$$\begin{aligned} \psi_{mRNA} =\left( \frac{0.205 g RNA}{g DW} \right)\times\left( \frac{0.04 g mRNA}{g RNA} \right)\approx0.8 \% \#\left( S1.9 \right) \end{aligned}$$

##### 4. Mass fraction of (all) proteins and ribosomes

Similar to above (2), determining the total *mass* fraction of proteins and ribosomes in the cell, $\psi_{proteins+ribosomes}$, amounts to simply determining the sum of mass fractions of proteins and rRNA, $\psi_{proteins}$ and $\psi_{rRNA}$ respectively:

$$\begin{aligned} \psi_{proteins+ribosomes}=\psi_{proteins}+\psi_{rRNA} \#\left( S1.10 \right) \end{aligned}$$

Recalling that $\psi_{rRNA} = f_{rRNA}\times\psi_{RNA}$, and replacing all parameters with the values from Table S1, we obtain:

$$\begin{aligned} \psi_{proteins+ribosomes}=\left( 0.55 \frac{g protein}{g DW} \right)+\left( 0.86\frac{g rRNA}{g RNA} \right)\times\left( 0.205\frac{g RNA}{g DW} \right)\approx73\% \#\left( S1.11 \right) \end{aligned}$$

##### 5. Mass fraction of ribosomes

Differently from the previous section, we are interested in determining the mass fraction of *only* ribosomes, $\psi_{ribosomes}$. For this, we need to estimate the mass fraction of ribosomal proteins, $\psi_{rProteins}$, as well the mass fraction of rRNA, $\psi_{rRNA}$:

$$\begin{aligned} \psi_{ribosomes}=\psi_{rProteins}+\psi_{rRNA} \#\left( S1.12 \right) \end{aligned}$$

where the mass fraction of ribosomal proteins is given by:

$$\begin{aligned} \psi_{rProteins}=f_{rProtein}\times\psi_{protein} \#\left( S1.13 \right) \end{aligned}$$

with $f_{rProtein}$ being the mass fraction of all proteins that are ribosomal proteins, and $\psi_{protein}$ the mass fraction of all proteins in the cell.

By replacing all parameters with the values from Table S1:

$$\begin{aligned} \psi_{ribosomes}=\left( 0.25\frac{g rProtein}{g protein} \right)\times\left( 0.55 \frac{g protein}{g DW} \right)+\left( 0.86\frac{g rRNA}{g RNA} \right)\times\left( 0.205\frac{g RNA}{g DW} \right) \approx31\% \#\left( S1.14 \right) \end{aligned}$$

### Supplemental Note 2 – DNA degradation under various conditions

To validate that the induction of ISce-1 expression with L-arabinose caused the expected DNA degradation, we incubated cells with DAPI (at a final concentration of 1 µg/mL, for 15 min, and at 37 °C) two hours after addition of the inducer. Since this dye becomes fluorescent when intercalated between two strands of DNA (56), the presence of fluorescence signal indicates that cells contain DNA. Thus, when we added DMSO to the cultures (either at the same time, or one hour after the addition of L-arabinose) most cells did not display any fluorescence in the appropriate imaging channel (λ_excitation_= 365 nm; Fig. S6A, left column), indicating that DNA was degraded.

In contrast, when we added rifampicin simultaneously with L-arabinose, most cells had a strong fluorescence signal (Fig. S6A, top right), suggesting they had not undergone DNA degradation. This observation is explained by the fact that rifampicin blocks all cellular transcriptional activity, including that of the *ISce-1* gene, and thereby prevents the expression of the DNA-cutting ISce-1 enzyme. Consistent with this view, when we delayed the addition of rifampicin until one hour after the addition of L-arabinose (thus allowing for an initial period during which the enzyme can be expressed), we observed that cells underwent DNA degradation similar to that seen in the DMSO-controls (Fig. S6A, bottom right).

Performing single particle tracking measurements on the four conditions described above allowed us to support two of the points made in the main text. First, we could reproduce the observation that diffusion of the particle is higher in cells devoid of DNA than in those still containing this macromolecule (Fig. S6B; *p* = 8.9 x 10^-21^, one-way ANOVA comparing all four distributions shown; see also Fig. 3B). Furthermore, we observed that in the condition where rifampicin was added one hour after L-arabinose (a treatment designed to enforce mRNA degradation after the onset of DNA degradation), the average diffusion coefficient was indistinguishable from that obtained under the conditions of DNA-degradation (Fig. S6B, bottom). This observation suggests that mRNA was absent in those conditions where DNA degradation was obtained by addition of L-arabinose and DMSO. Importantly, this validates our assumption (see Fig. 1 and main text) that DNA-free cells are also mRNA-free.

### Supplemental Note 3 – Mathematical model of crowding

To relate the observed changes in the diffusion coefficients upon various treatments to changes in the levels of intracellular crowding, we used the model of Mendoza & Santamaría-Holek (2009), Eq. S2.1, a generalization of Einstein’s equation for the viscosity of a dilute suspension of *hard spheres* to a regime of arbitrary crowder concentrations:

$$\begin{aligned} \eta=\eta_{o}\left( 1-\Phi_{eff} \right)^{-\frac{5}{2}} \#\left( S2.1 \right) \end{aligned}$$

where $\eta$ and $\eta_{o}$ are the viscosities of the crowded suspension and the solvent, respectively, and $\Phi_{eff}$ is the effective filling fraction. At low crowder concentrations, $\Phi_{eff}$ tends to $\Phi$, the volumetric fraction of crowders. Given the presence of long polymeric molecules, such as mRNA or DNA, in the cytoplasm, a model that assumes spherical crowder geometry is clearly an unrealistic approximation. Nevertheless, the estimation of excluded volume due to polymers in terms of an *equivalent* amount of spherical crowders is still insightful by allowing us to set an upper bound on the true excluded volume experienced by a tracer particle.

Since the diffusion coefficient is inversely proportional to the viscosity of the medium, Eq. S2.1 allows us to derive the following relation between the diffusion coefficient in solvent only and in the crowded suspension:

$$\begin{aligned} \frac{D_{o}}{D}=\left( 1-\Phi_{eff} \right)^{-\frac{5}{2}} \#\left( S2.2 \right) \end{aligned}$$

Converting both sides of Eq. S2.2 to log_10_-scale, we arrive at:

$$\begin{aligned} \log_{10} D_{o}-\log_{10} D= -\frac{5}{2}\log_{10} \left( 1-\Phi_{eff} \right) \#\left( S2.3 \right) \end{aligned}$$

Using Eq. S2.3, we can now generalize to a scenario where diffusion is compared between two conditions where crowders are present:

$$\begin{aligned} {\Delta\log_{10} D\left. \right|_{cond1\to cond2}= log}_{10} D_{cond1}-\log_{10} D_{cond2}= -\frac{5}{2}\log_{10} \left( \frac{1-\Phi_{cond2}}{1- \Phi_{cond1}} \right) \#\left( S2.4 \right) \end{aligned}$$

In accordance with Fig. 1, we can relate each experimental condition with the macromolecular crowders present. In particular, in the control condition, we have ${\Phi_{DMSO}= \Phi}_{mRNA+DNA+others}$; under rifampicin treatment, ${\Phi_{Rifampicin}= \Phi}_{DNA+others}$; and under induction of ISce-1 expression with L-arabinose, ${\Phi_{L-arabinose}= \Phi}_{others}$.

If we then assume that the total excluded volume of a mixture of crowders can be decomposed in the sum of the excluded volume of its constituents, that is:

$\begin{aligned} \Phi_{i+j+k+\ldots}=\Phi_{i}+\Phi_{j}+\Phi_{k}+\ldots\#\left( S2.5 \right) \end{aligned}$

we can derive, from Eq. S2.4, the following expressions to determine the excluded volume due to each macromolecular crowder in the cell from our experimental data (log_10_*D* values):

$$\begin{aligned} \Phi_{others}= 1-{10}^{-\frac{2}{5} \left( {\Delta log}_{10} {D|}_{o\to L-arabinose} \right)} \#(S2.6.a) \end{aligned}$$

$$\begin{aligned} \Phi_{DNA}=\left( 1-\Phi_{\mathrm{others}} \right)\left( 1-{10}^{-\frac{2}{5} \left( \Delta\log_{10} {D|}_{L-arabinose\to Rifampicin} \right)} \right)\#\left( S2.6.b \right) \end{aligned}$$

$$\begin{aligned} \Phi_{mRNA}=\left( 1-\Phi_{\mathrm{others}}-\Phi_{DNA} \right)\left( 1-{10}^{-\frac{2}{5} \left( \Delta\log_{10} {D|}_{Rifampicin\to DMSO} \right)} \right)\#(S2.6.c) \end{aligned}$$

### Supplemental Table

Table S1 – Parameters obtained from the literature. Compilation of values describing the macromolecular composition of bacterial cells and properties of some macromolecules

| Parameter | Name | Value | Reference |
| --- | --- | --- | --- |
| $\boldsymbol{f}_{\boldsymbol{mRNA}}$ | (Mass) fraction of RNA that is mRNA | 4% | (57) |
| $\boldsymbol{f}_{\boldsymbol{rRNA}}$ | (Mass) fraction of RNA that is rRNA | 86% | (58) |
| $\boldsymbol{f}_{\boldsymbol{rProtein}}$ | (Mass) fraction of proteins that are ribosomal proteins | ~25% | (59) |
| $\boldsymbol{\psi}_{\boldsymbol{protein}}$ | (Mass) fraction of cellular dry weight that is protein | 55.0% | (60) |
| $\boldsymbol{\psi}_{\boldsymbol{RNA}}$ | (Mass) fraction of cellular dry weight that is RNA | 20.5% | (60) |
| $\boldsymbol{\nu}_{\boldsymbol{RNA}}$ | Partial specific volume of RNA | 0.569 mL/g | (61) |
| $\boldsymbol{\nu}_{\boldsymbol{protein}}$ | Partial specific volume of proteins | 0.728 mL/g | (61, 62) |
| $\boldsymbol{c}_{\boldsymbol{RNA}}$ | Cellular RNA concentration | 75 – 120 mg/mL | (1) |
| $\boldsymbol{\rho}_{\boldsymbol{cell}}$ | Cellular dry density | 0.3 gDW/mL | (63) |

### Supplemental Figures (S1 – S8)

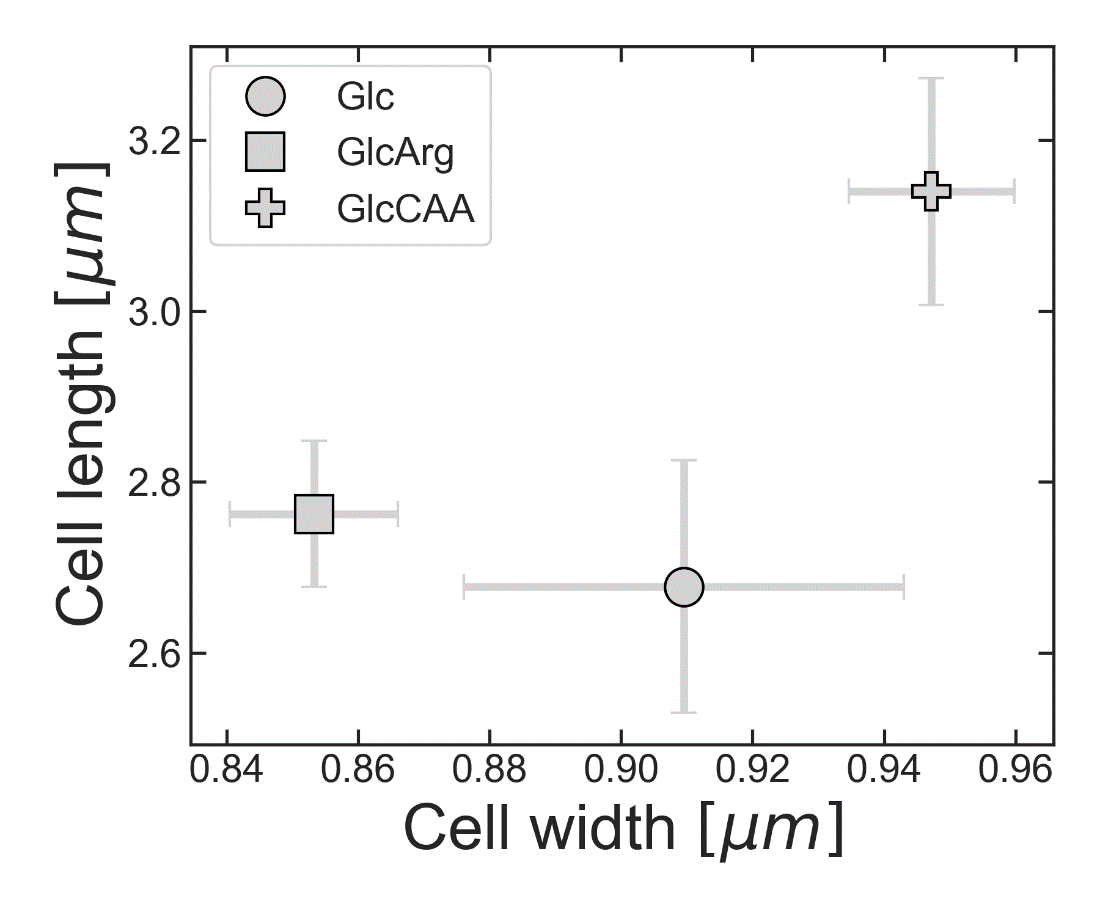

Figure S1 – Morphology of cells under different growth conditions.

Length and width of *E. coli* LY177 cells grown in minimal medium with glucose (Glc), glucose and L-arginine (GlcArg), or glucose and casamino acids (GlcCAA). The values refer exclusively to cells without visible constriction rings. Each marker shows the mean ± standard deviation of n = 15 experimental replicates, where the value of each experimental replicate was obtained by taking the mean across individual cells.

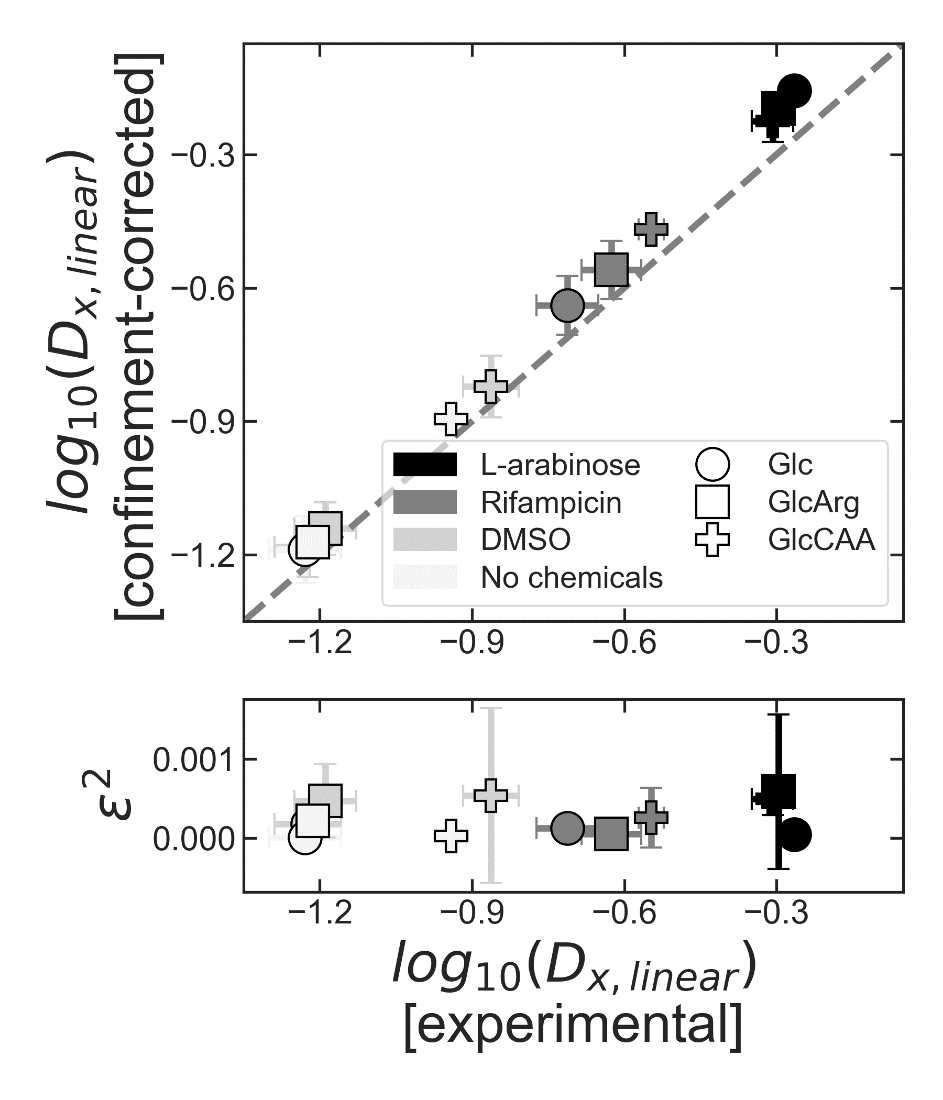

Figure S2 – Diffusion coefficients corrected for the effect of confinement are similar to the uncorrected values.

(top) Diffusion coefficients corrected for the effect of confinement by the cell membrane (y-values; data displayed in Fig. 3A and B) are plotted against the experimentally determined diffusion coefficients for the same condition (x-values). The corrected values were obtained using the iterative procedure outlined in Fig. 2C. Note that the higher the diffusion coefficient of the confined particles, the larger the discrepancy between the experimental (i.e., affected by confinement) and the real (i.e., not affected by confinement) values of that parameter. The dashed line represents the diagonal, y = x. (bottom) Average values of the squared residuals, *ε^2^*, a measure for how well the estimated diffusion coefficients corrected for the effect of confinement fit the experimental values. Each marker shows the mean ± standard deviation of n ≥ 3 experimental replicates.

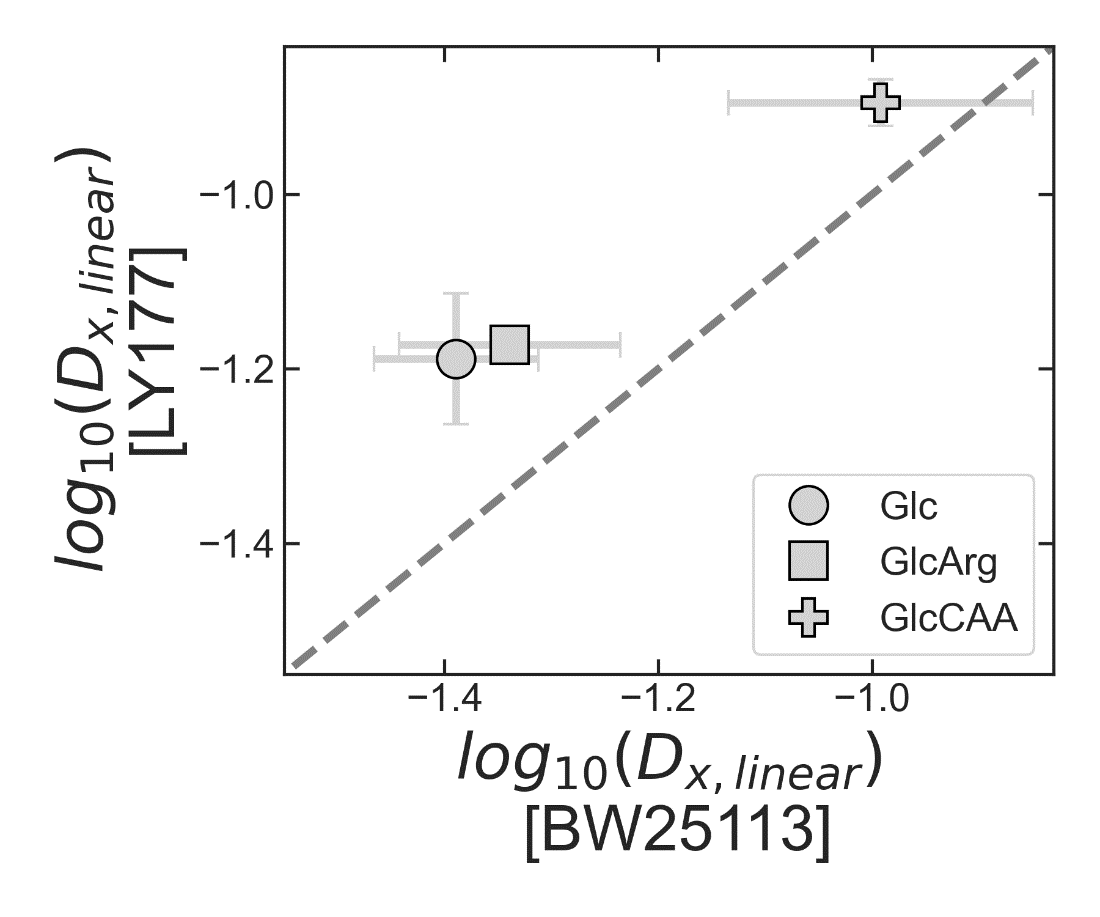

Figure S3 – Diffusion coefficients measured in LY177 and BW25113 strains are different but show the same relative trend.

Confinement-corrected diffusion coefficients measured in LY177 and BW25113 *E. coli* strains grown in different media (Glc, GlcArg or GlcCAA; no other chemicals added) are plotted against each other. The dashed line represents the diagonal, y = x. Each marker shows the mean ± standard deviation of n ≥ 3 experimental replicates.

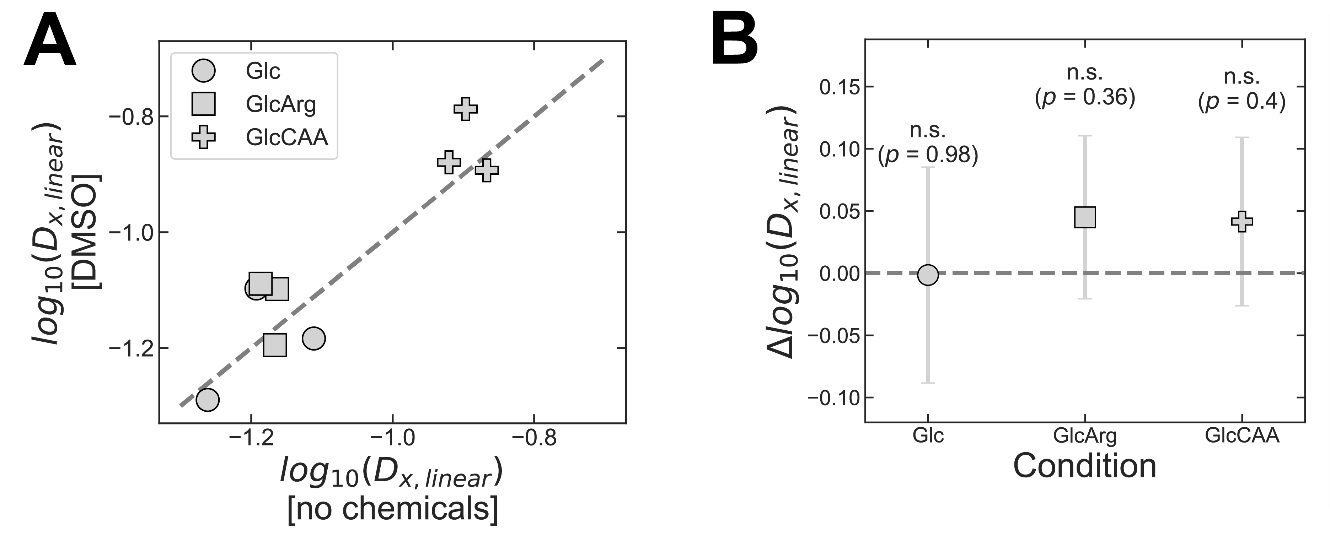

Figure S4 – Addition of DMSO does not affect the estimated diffusion coefficients.

(**A**) Confinement-corrected diffusion coefficients measured in the LY177 strain, grown in different media (Glc, GlcArg or GlcCAA), in n = 3 independent experiments. After a common preculturing period, the cultures were split, and single particle tracking was done in the presence or absence of DMSO ("DMSO" and "no chemicals", respectively). The dashed line represents the diagonal, y = x.

(**B**) Difference between the confinement-corrected log_10_*D* values under the two treatments ("DMSO" and "no chemicals"). The values of ∆log_10_*D* were calculated for each experimental replicate (i.e., difference between the y- and x- values shown in (**A**)), and the mean ± standard deviation across the n = 3 replicates is shown by the markers. Given the high *p*-values (two-sided t-test for paired samples), the values of ∆log_10_*D* are not statistically different from 0 ("n.s.").

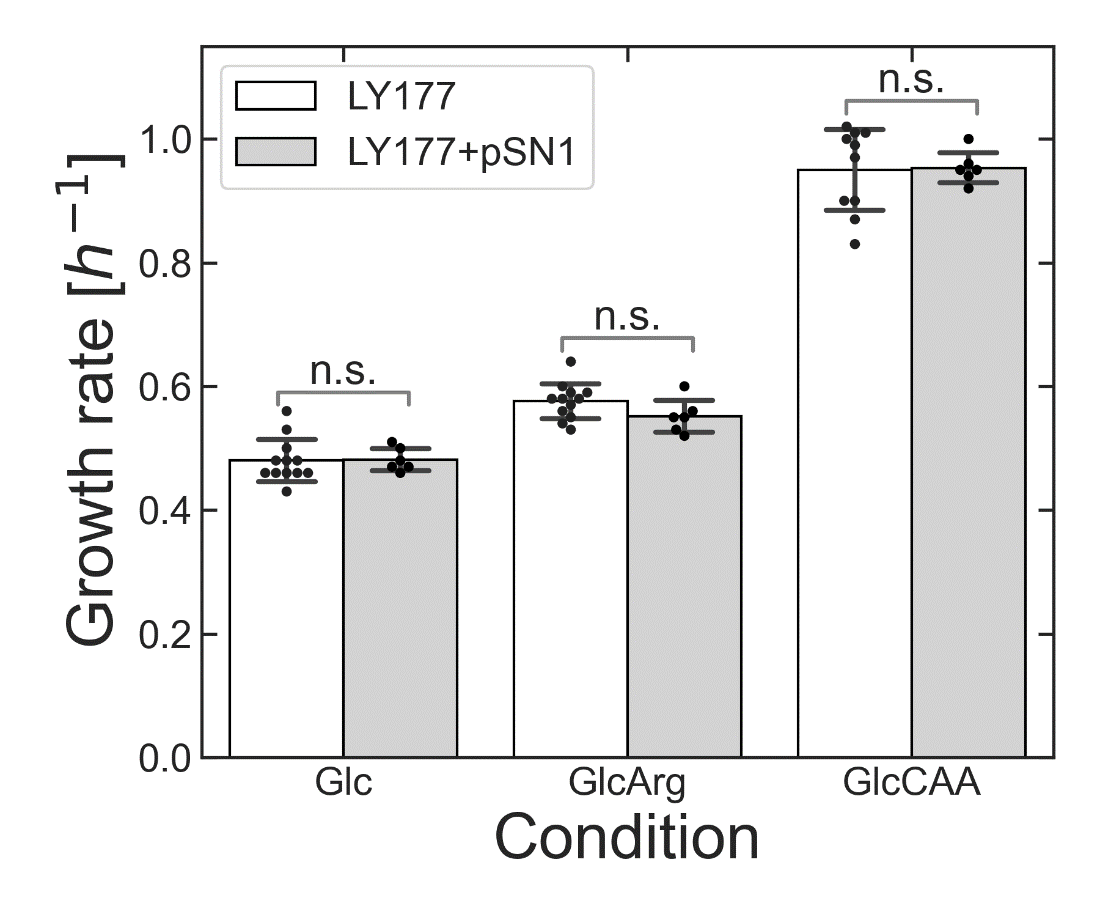

Figure S5 – Strains LY177 and LY177+pSN1 have similar growth rates.

Strains LY177 and LY177+pSN1 were grown in different media (Glc, GlcArg or GlcCAA), and their growth rates were determined prior to any of the treatments described in the main text. The values from independent experiments are plotted as black dots and their mean values are given by the height of the respective bars. For all conditions, the difference between the growth rates of the two strains is statistically not significant, “n.s.” (*p* ≥ 0.11, two-sided t-test for independent samples).

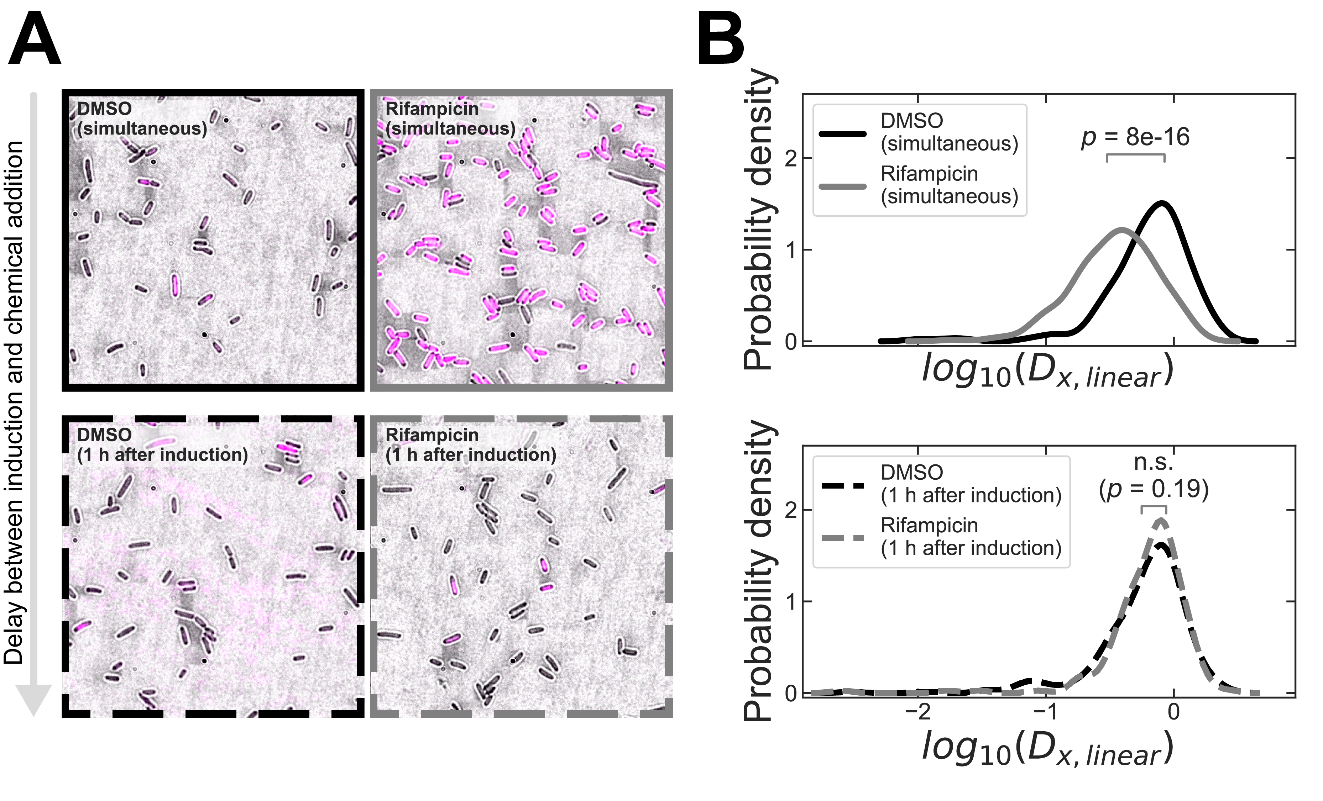

Figure S6 – DNA-degradation results in increased particle diffusion coefficients.

(**A**) Overlaid brightfield and fluorescence channel (λ_excitation_ = 365 nm; λ_emission_ = 415-455 nm) images of DNA-stained LY177+pSN1 cells. Cells were induced with L-arabinose, for expression of ISce-1, and were additionally exposed to: DMSO, added at the time of induction (top left); rifampicin, added at the time of induction (top right); DMSO, added one hour after induction (bottom left); rifampicin, added one hour after induction (bottom right). In all cases, cells were stained two hours after induction with L-arabinose, by incubation with DAPI for 15 min, at 37 °C. The fluorescence channel has the same scale in all four images. The images were obtained with 150x magnification.

(**B**) Distribution of diffusion coefficients measured in LY177+pSN1 cells, not corrected for the effect of confinement, two hours after induction with L-arabinose. Cells were grown in minimal medium with glucose. Either DMSO or rifampicin were added to the cultures. These two chemicals were added either at the same time as (top), or one hour after addition of L-arabinose (bottom). As shown in (**A**), cells in all conditions but “Rifampicin (simultaneous)” underwent DNA-degradation. Lines are color- and style-coded to match the frames in (**A**). Equality of the means of the distributions was assessed by a one-way ANOVA statistical test (*p*-values shown in each plot).

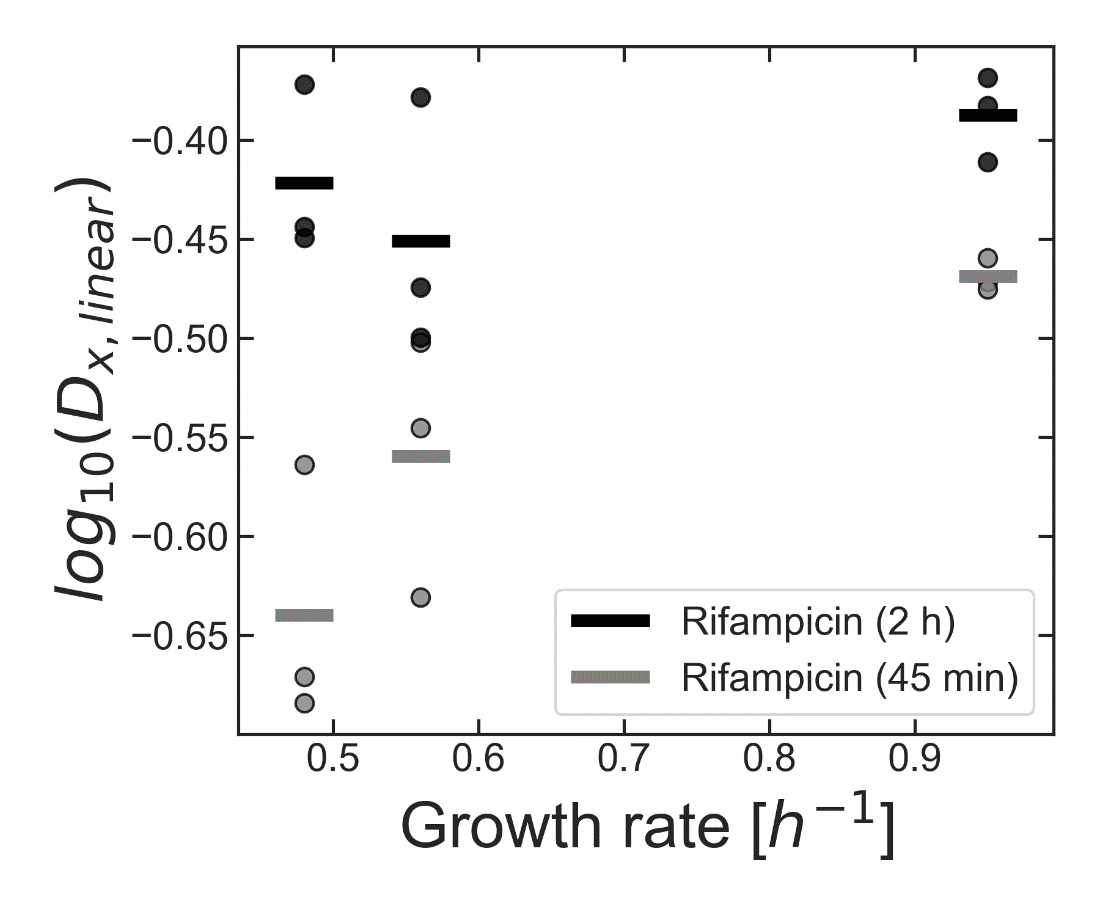

Figure S7 – Duration of the exposure to rifampicin affects the diffusion coefficients.

Confinement-corrected diffusion coefficients were measured in LY177 cells exposed to rifampicin for 45 min, and in LY177+pSN1 exposed to rifampicin and L-arabinose for 2 hours. Values in the latter case are consistently higher than in the former. While time-dependent diffusion coefficients upon exposure to rifampicin have been previously reported for chromosomal loci (44) and cytoplasmic particles (64), these previous findings rather indicate that diffusion coefficients should plateau at ~30 min after the onset of the treatment. While it remains unclear how our observations can be reconciled with the observations from those two studies, our results could be explained in light of the recent findings of Hamouche *et al* (2021). These authors report that not only mRNA, but also rRNA is degraded when cells are exposed to rifampicin. Thus, it is conceivable that even ribosomes undergo partial degradation during the period between 45 min and 2 h after exposure to the chemical. The resulting reduction in crowding could thereby lead to the observed increase in the diffusion coefficient. The data referring to "Rifampicin (45 min)" is the same as the "Rifampicin" set in Fig. 3B. The horizontal bars represent the average values obtained in n = 3 experiments (markers).

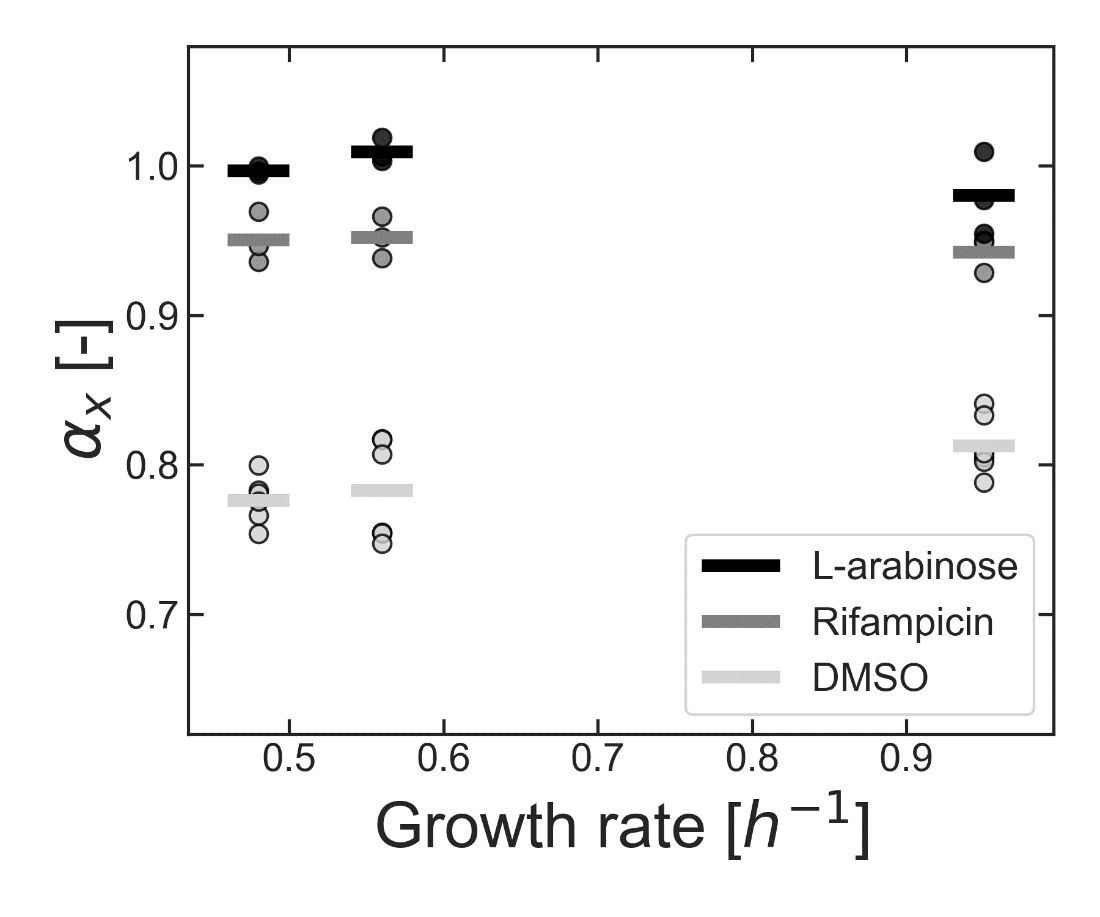

Figure S8 – Anomalous diffusion exponents show changes in the diffusive regime across different experimental treatments.

The anomalous diffusion exponents, *α*, obtained in cells grown in different media (Glc, GlcArg, GlcCAA), and subjected to different experimental treatments (same as in Fig. 3B). The values of *α* were obtained by fitting particle mean squared displacements, *MSD*, to the equation $MSD = 2nD\tau^{\alpha}$, instead of Eq. 1 (which was used elsewhere in this work). The subscript "x" indicates that the data refers to motion along the long cell axis. The horizontal bars represent the average values obtained in n ≥ 3 experiments (markers).
